# Model-based Evaluation of Connexin Hemichannel Permeability

**DOI:** 10.1101/2025.06.19.660562

**Authors:** Tadas Kraujalis, Lukas Kersys, Andrius Krisciunas, Dalia Calneryte, Auguste Cicinskaite, Lina Kraujaliene, Vytas K. Verselis, Mindaugas Snipas

## Abstract

Connexin (Cx) hemichannels represent a growing facet of Cx research. Hemichannels are hexamers of Cx subunits that are historically thought of as substrates of intercellular gap junction channels formed when two hemichannels from neighboring cells dock. Hemichannels, however, can function in the absence of docking and have been shown to play important roles in transmembrane signaling. There are 21 different Cx isoforms in humans and a host of disease-causing mutations have been identified. Although Cxs constitute large-pore channels, they can differ substantially in their conductance and permeability characteristics, thereby affecting the nature of signals that are transmitted. In this study, we present a methodology for quantifying and comparing the permeabilities of hemichannels formed by different Cx isoforms using a combination of fluorescence imaging, electrophysiological recordings and mathematical modeling. Fluorescence imaging, coupled with mathematical modeling based on Fick’s law and/or the Goldman-Hodgkin-Katz current equation, enables estimation of tracer diffusion rates. These data are integrated with independently obtained electrophysiological measurements of hemichannel activity into a unified statistical model based on the likelihood ratio test. Simulation-based analyses demonstrate that our approach can reliably detect ∼2-fold differences in hemichannel permeability using datasets of moderate size. Crucially, this approach requires only a minimal amount of time-intensive electrophysiological recording, leveraging higher-throughout fluorescence measurements, which can be further streamlined using AI-based tools for automated cell detection and data extraction. We apply this method to compare the permeability of hemichannels formed by wild-type Cx26 and a pore-lining variant, Cx26^*^A49E. Our results show a significant increase in DAPI permeability in Cx26*A49E hemichannels, consistent with previous findings. This methodology can be extended to assess the permeabilities of other large-pore channels.

## Introduction

Connexin (Cx) proteins are the substrates of large-pore channels that can function both in transmembrane signaling and direct intercellular signaling. This dual function comes from the ability of Cxs to assemble and operate in two configurations, as gap junction (GJ) channels that function in direct intercellular signaling between cell cytoplasms, formed by the docking of two hexameric channels, termed hemichannels, one from each of two neighboring cell membranes and as undocked hemichannels that function in the plasma membrane much like conventional ion channels. There are 21 different Cxs in the human genome that show overlapping, but tissue-specific patterns of expression. Because of their generally large pore size, Cx channels facilitate electrical as well as chemical signaling in a variety of tissues.

Hemichannels likely transmit a variety of signaling molecules and have garnered increased attention in transmembrane signaling, with reports that they play both physiological and pathological roles in varied processes such as calcium wave propagation [1], ATP release [2; 3], response to metabolic stress [4; 5], and inflammation [6; 7]. In addition, aberrant hemichannel function has been associated with Cx mutations in human disease, most notably in syndromic, sensorineural deafness, in which mutations in Cx26 and Cx30 cause profound hearing loss accompanied by cutaneous manifestations, both infectious and neoplastic, with phenotypes that vary based on the specific mutation (revs. in [8; 9]). Increased hemichannel open probability and/or altered permeability has been reported in syndromic deafness, which leads to cellular dysfunction through enhanced passage of harmful molecules and/or the restricted passage of essential signaling molecules [10; 11; 12; 13; 14].

Hemichannel permeability is of particular interest as emerging Cx structures show substantially different electrostatic landscapes within the pores, which is not surprising given the >10 fold range of unitary conductance reported for GJs composed of different Cx isoforms (rev. in [15]). Also, disease-causing Cx mutations that cause syndromic deafness often occur in the NT and E1 domains that constitute the bulk of the pore and, thus, likely alter hemichannel permeability. While the permeabilities of Cx hemichannels to ions can be assessed from conductance measurements using conventional patch clamp techniques, permeabilities to larger molecules requires different approaches. Molecular permeability is typically assessed using uptake or leakage assays of fluorescent or radioactive tracers. Fluorescent dyes of various sizes and charges can provide permeability profiles that are proxies for natural permeants and uptake and/or release assays are widely used to assess Cx hemichannel permeability.

To quantify hemichannel permeability requires explicitly quantifying hemichannel numbers in uptake/release assays, which can be obtained by a combination of whole-cell and single channel recording. This combination provides the mean number of hemichannels that are open at any given time. The difficulty is that the introduction of a patch electrode in a whole-cell configuration disrupts the measurement of intracellular fluorescence through dialysis of cellular contents. As a solution, studies have used large *Xenopus* oocytes to express Cxs and measure tracer flux in conjunction with membrane conductance using a two-electrode voltage clamp and sharp electrodes [16; 17; 18]. However, oocytes have excessively large volumes, thereby requiring long observation times, typically exceeding 60 minutes to observe substantive fluorescence changes making it difficult to maintain a stable recording. In addition, *Xenopus* oocytes are pigmented which hampers fluorescence detection, requiring arduous additional preparation steps involving centrifugation and imaging configurations to improve detection [19]. And finally, measurements of tracer flux under conditions of voltage clamp that impose substantial transjunctional voltages to monitor conductance, have not generally taken into account the influence of the transmembrane field.

In this study, we propose a method for comparing Cx hemichannel permeabilities in mammalian cells that integrates electrophysiological recording, fluorescent imaging, and mathematical/statistical modeling. An advantage of this methodology is that electrophysiological recordings and tracer uptake/release assays are performed separately and then compared using a statistical model, making it technically less challenging than approaches that employs simultaneous electrophysiological recordings and fluorescence imaging in the same cells. Specifically, total and unitary hemichannel conductances are measured to provide estimates of the average number of open hemichannels in a given cell population. In separate cells, fluorescence imaging data from tracer uptake/release assays are combined with mathematical modeling to assess the diffusion rate, which is proportional to the permeability of a single hemichannel. To analyze fluorescence imaging data, we employ machine learning-based cell detection and segmentation, which automates the extraction of fluorescence intensity changes over time and enables cell volume estimation, which is relevant for diffusion rate calculations. The independently obtained conductance and diffusion rate data are then integrated into a statistical model based on likelihood ratio test, allowing determination of whether hemichannel permeability differs significantly, with associated p-values. Our simulation-based analysis indicates that this method can detect permeability differences as small as ∼2-fold with moderate sample sizes.

We demonstrate our methodology using cell cultures expressing WT Cx26 and a Cx26^*^A49E variant, in which Glu at position 49 is replaced with Ala. Previous studies [20; 21; 22] identified this residue as a key determinant of permeability difference between Cx26 and Cx30 hemichannels. Our results show that Cx26^*^A49E hemichannels exhibit significantly higher permeability to the positively charged DAPI dye compared to WT channels. This finding aligns with prior studies, further supporting the role of the negatively charged Glu at position 49 as a molecular determinant of Cx26 hemichannel permeability as it relates to the charge of the permeant.

Overall, our study demonstrates that the proposed methodology effectively compares the permeability of different Cx hemichannels. In principle, this approach could be extended to other large-pore channels, such as pannexins, calcium homeostasis regulators (CALHMs), and volume-regulated anion channels (VRACs). A major advantage of this method is that electrophysiological and fluorescent imaging data can be collected separately, allowing for greater experimental flexibility. Moreover, our simulations suggest that increasing sample sizes for either conductance measurements or fluorescence imaging enhances statistical power to a similar extent. Thus, our approach facilitates high-throughput fluorescent imaging, enabling efficient permeability comparisons and reducing the demand for time-intensive electrophysiological recordings.

## Methods

### Cell Cultures

HeLa mammalian cell lines (ATCC CCL-2) were used for Cx expression. Both wild-type Cx26 and its mutant Cx26^*^A49E were fused with msfGFP to enable identification of transfected cells. Cell cultures were maintained in Dulbecco’s Modified Eagle Media (DMEM; Sigma, USA) supplemented with 10 % fetal bovine serum (Sigma, USA) and 1% penicillin-streptomycin solution (10,000 units penicillin and 10 mg/ml streptomycin; Sigma, USA). Cells were incubated under standard conditions of 5% CO_2_ at 37 °C. For transient transfection, HeLa cells were transfected on the second day after seeding onto coverslips in Petri dishes using either Lipofectamine 2000 (Invitrogen, USA) or jetPrime (Polyplus Transfection, France) reagents. Experimental procedures were conducted 48 hours post-transfection.

### Electrophysiological Recordings

For electrophysiological and fluorescence recordings, HeLa cells grown on glass coverslips were transferred to an experimental chamber mounted on the stage of an Olympus IX70 inverted microscope (Olympus, Japan) equipped with a fluorescence imaging system. Extracellular solution was applied to the bath near the cell using a gravity-driven perfusion system at a flow rate of approximately 2 mL/min. All experiments were conducted at room temperature. Single whole-cell patch-clamp methods were used to record currents from hemichannels using EPC-8 patch-clamp amplifiers (HEKA Elektronik, Germany). Currents were digitized at a sampling rate of 5 kHz and low-pass filtered at 1 kHz. Hemichannel recordings were obtained from voltage steps and ramps applied to single, isolated cells clamped in a whole-cell configuration, which exhibited fluorescent dots in the cell area. Resting membrane potential was measured using a whole-cell patch-clamp set to a current-clamp mode (with 0 current) during the first five seconds after obtaining the seal.

Patch pipettes were fabricated from borosilicate glass tubes with filaments (BF 150-86-10, Sutter Instrument Co., USA) using a P-97 micropipette puller (Sutter Instrument Co., USA) to achieve a resistance between 2–3 MΩ, minimizing the effects of series resistance on current measurements. The intracellular solution contained (in mM): 130 CsCl, 10 NaAsp, 0.26 CaCl_2_, 5 HEPES, 2 BAPTA, and 1 MgCl_2_, with a pH of 7.7. The divalent-free extracellular solution contained (in mM): 140 NaCl, 4 KCl, 2 CsCl, 1 BaCl_2_, 5 HEPES, 5 glucose, and 2 pyruvate, with a pH of 7.8. For the measurements of resting membrane potentials, CsCl was replaced with KCl.

Data were acquired using AT-MIO-16X D/A boards from National Instruments and analyzed with custom acquisition and analysis software (VTDaq, NexusWiz, written by E. Brady Trexler, Gotham Scientific, Hasbrouck Heights, NJ, USA).

### AI-based Cell Detection System for Fluorescence Imaging Experiments

#### Fluorescent imaging experiments

HeLa cells were grown on glass coverslips, placed in an experimental chamber containing an extracellular solution, and imaged using an Olympus IX70 inverted microscope (Olympus, Japan) equipped with a fluorescence imaging system. Brightfield and fluorescence images were captured using an ORCA digital camera (Hamamatsu, Japan) and processed with UltraVIEW imaging software (PerkinElmer Life Sciences, Boston, MA, USA). Images were acquired with a fixed magnification to ensure consistency across the dataset.

For DAPI (4’,6-diamidino-2-phenylindole) uptake experiments, HeLa cells transfected with either wild-type (WT) Cx26 or Cx26^*^A49E were seeded on glass coverslips and placed on the same setup as described above. The cells were bathed in a divalent-free extracellular solution for 5 minutes to open hemichannels. Subsequently, perfusion with an external solution containing 5 μM DAPI was initiated simultaneously with the start of the recording and maintained at a constant flow rate to ensure a steady supply of the dye throughout the recording period. Bright-field and fluorescence images with EGFP filters were captured for a field of cells prior to DAPI uptake. To monitor DAPI uptake, changes in intracellular fluorescence intensities were recorded every 15 seconds (100 ms exposure time) for the 30-minute recording duration. Appropriate excitation and emission filters were used to visualize DAPI and tagged Cx variants.

#### Dataset and Annotation

Cell boundary detection was performed using microscopy images of HeLa cells, as detailed in the cell imaging section. Annotation of cell boundaries was done using the online annotation tool provided by makesense.ai. 72 images containing 1618 cells was used for training and 32 images containing 139 cells for validation. Each image had a resolution of 1344 × 1024 pixels, with a pixel size of 0.1075 um.

#### Model Architecture and Training

Automated segmentation of cell boundaries was executed using the Detectron2 library, chosen due to its superior segmentation capabilities compared to other advanced methods. The Mask R-CNN architecture with a ResNet-50 backbone combined with a Feature Pyramid Network was applied using the mask_rcnn_R_50_FPN_3x.yaml configuration. The model was trained for up to 4500 iterations with an initial learning rate set to 0.00025. To improve detection accuracy, a positive Intersection-over-Union (IoU) threshold of 0.5 was used to identify true positives, and a Non-Maximum Suppression threshold of 0.75 was applied to eliminate overlapping detections. Predicted segmentations covering less than 0.3% of the total image area were also excluded to reduce noise. As the dataset contains only a single cell class, the number of classes was set to 1. Training and validation datasets were formatted following the COCO annotation style, and training was carried out on an NVIDIA A100-PCIE-40GB GPU.

#### Post-processing of Segmentation Results

Post-processing was performed after segmentation to enhance accuracy and consistency. Multi-polygon filtering ensured that each cell was represented by a single continuous segment, retaining only the largest polygon when multiple polygons were produced for one cell. Overlapping segmentations were evaluated using the intersection coefficient (IC), defined as the ratio of the overlapping area to the smaller of the two overlapping areas. Segmentations with an IC greater than 0.5 were resolved by removing the detection with the lower confidence score.

#### Performance Evaluation

After validation, the model achieved a precision of 75.45% and a recall of 90.65%. Following post-processing, the overall precision improved to 93.94% on the validation dataset, indicating about 6% false-positive detections, whereas recall reduced to 89.21% after filtering from 90.65% before filtering – meaning about 10% of cells missed detection completely. After filtering, the overall accuracy increased from 70.00% to 84.35%. Additionally, the segmentation area accuracy, measured by the IoU between predicted boundaries and ground truth, showed an intersection error of less than 1%.

#### Computations and Statistical Analysis

Mathematical and statistical models were implemented in MATLAB.

## Results

### 1. Mathematical model for tracer molecule uptake assays

#### 1.1 Changes in concentration of tracer molecule

We begin with a general mathematical model describing the flux of tracer molecules across a cell membrane mediated by permeable pores. The rate of concentration change between the cell cytoplasm and extracellular space can be described using a two-compartment model, where diffusion is governed by Fick’s law. Specifically, the rate of concentration change is proportional to the concentration gradient between the inside and outside of the cell.

Denoting the concentrations of tracer molecules inside and outside the cell at time *t* as *C*_*in*_(*t*) and *C*_*out*_(*t*), respectively, and defining the volumes of the cell and bathing medium as *Vol*_*in*_ and *Vol*_*out*_, respectively, the rate of concentration change is governed by the following system of ordinary differential equations (ODEs):

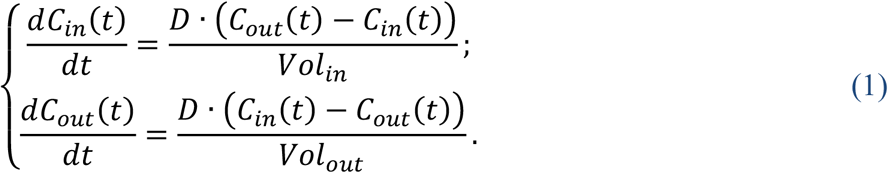

Here variable *D* represents the diffusion rate constant, which has the dimensions of volume per unit time. The diffusion rate constant can also be expressed as the product of the single-channel permeability to the permeant molecule *P* (which also has the dimensions of volume per unit time), the number of channels *n*, and channel open probability *p*_*o*_:

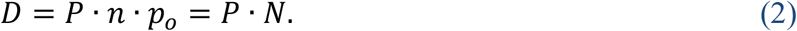

Here, *N* = *n*·*p*_*o*_ denotes number of open channels. Similarly, the macroscopic channel conductance to ions, *g*, is the product of the single-channel conductance *γ*, the number of channels *n*, and channel open probability *p*_*o*_:

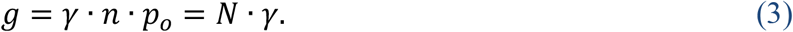

This formulation allows the permeability of a single open channel, *P*, to be evaluated from the relationship

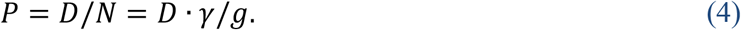

Since the volume of the bathing medium is significantly larger than the volume of the cell (*Vol*_*out*_ >> *Vol*_*in*_) the rate of change in *C*_*out*_, *dC*_*out*_(*t*)/*dt*, is negligible and can be approximated as zero. Thus, we can assume *C*_*out*_ to be constant, *C*_*out*_(*t*) = *C*_*out*_. This assumption simplifies the system of ODEs in Eq. (1) to a single ODE:

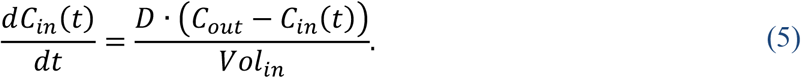

At the beginning of the tracer uptake assay experiment, the initial concentration of tracer molecules inside the cells is approximately zero, *C*_*in*_(0) ≈ 0. The solution of Eq. <u>(5)</u>is the given by:

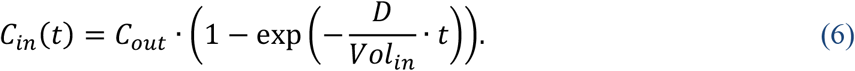

The expression based on Fick’s law-based Eq. (5) is valid when the tracer molecule is electrically neutral or when the membrane potential V_m_ is at or close to 0 mV. However, many tracer molecules are charged, and cells often maintain significant resting membrane potentials. In such cases, diffusion is also influenced by the electric field across the membrane.

To account for the effects of a transmembrane field, we incorporate the Goldman-Hodgkin-Katz (GHK) current equation, which describes the flux of charged particles under the influence of both concentration and voltage gradients. The modified rate equation becomes:

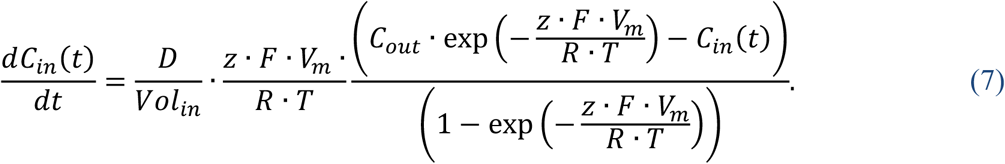

Here *z* is the valence of the tracer molecule, *F* is Faraday’s constant, *R* is the gas constant, and *T* is an absolute temperature in kelvins.

For convenience, we define a dimensionless potential parameter *μ*= (*z* · *F* ·*V*_*m*_)/(*R* ·*T*), the effective diffusion coefficient *D* ^*\**^ *=D* · *μ*/(1 − exp(− *μ*)), and the effective external concentration 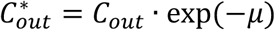. Using these definitions, Eq. (7) can be rewritten in a form analogous to Eq. (5)

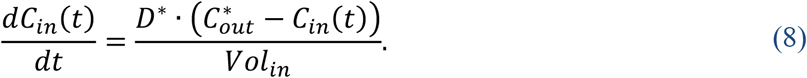

The solution of this equation is structurally similar to Eq. (6):

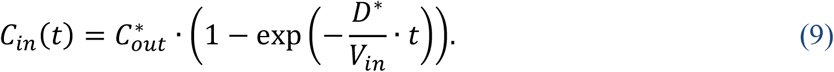

For uncharged tracer molecules (z = 0) or when the membrane potential is zero (V_m_ = 0 mV), the dimensionless parameter β = 0. By applying l’Hopital’s Rule, it can be shown that *β* /(1 − exp (− *β*)) approaches 1 when β approaches 0, reducing Eq. (7) to the original Fick’s law-based formulation in Eq. (5).

The ODEs in Eq. (1) can also be adapted to describe intracellular concentration changes during dye release assays. In this case, the concentration of the tracer molecule in the bathing medium remains effectively zero due to its much larger volume relative to the cell. Consequently, the dynamics of *C*_*in*_(*t*) can are governed by:

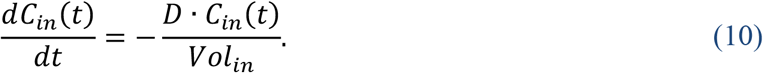

The solution to this equation describes an exponential decay of intracellular tracer concentration from the initial value *C*_*in*_(0) towards zero:

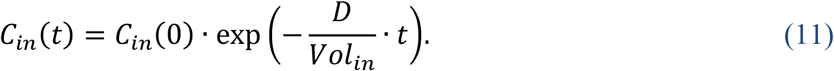

For charged tracer molecules, this model can be further extended to account for the influence of the resting membrane potential using the GHK formulations. Similar to the dye uptake model, this is achieved by replacing *D* in Eq. (11) with the effective diffusion coefficient *D* ^*^ =*D* · *μ* /(1 −exp (− *μ*)), where *β* incorporates the effects of the membrane potential as previously defined.

While our primary analysis presented in the following sections focuses on dye uptake assays, the same methodology can be readily adapted to describe dye release experiments.

#### 1.2 Relationship between changes in concentration and fluorescence intensity

Under experimental conditions, the concentration *C*(*t*) is assessed from the fluorescent intensity, *F*(*t*), which for many tracers depends linearly on the concentration with the ranges used [23; 24]. Thus, *F*(*t*), can be described by the following relationship:

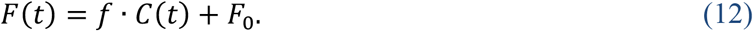

Here *f* is a constant denoting fluorescence per unit concentration, and *F*_0_ is the background fluorescence, representing the fluorescence in the absence of tracer molecules. Both *f* and *f*_0_ are measured in artificial units (*a. u*.) per unit of concentration.

Thus, in a tracer uptake assay, the increase in fluorescent intensity inside the cell, *F*_in_(*t*), is given by:

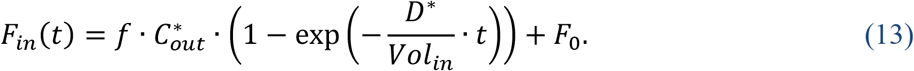

The background fluorescence can be measured at the beginning of the tracer uptake experiment, when *C*_*in*_(*t*) is equal to zero:

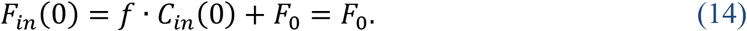

By subtracting the background fluorescence and normalizing to the initial fluorescence level, we obtain the following relationship:

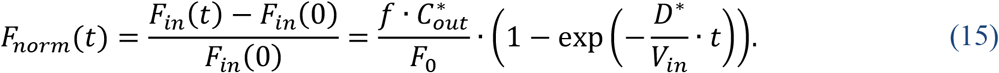

For tracer release assays described by Eq. (11), a very similar expression to Eq. (15) can be obtained by considering background-subtracted fluorescent recordings *F*_*in*_(0) – *F*_*in*_(*t*). More precisely, it has the following form:

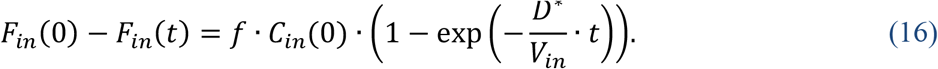

#### 1.3 Fitting mathematical model to experimental data

Equations (15) and (16) predict that the normalized and/or background-subtracted fluorescence intensity obtained in tracer uptake and release assays can be well described by an exponential function of the form

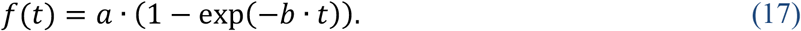

where parameter *a* corresponds to 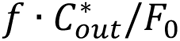, and *b* reflects the effective diffusion rate per unit cell volume, *D*^*\**^/*Vol*_*in*_. This function predicts the saturation of normalized fluorescence intensity as the concentration of tracer molecules inside the cell, *C*_*in*_(*t*), approaches that of the bathing medium, *C*_*out*_.

When the effective diffusion rate is not large, the time course of *F*_*norm*_(*t*) should not exhibit an immediate inflection toward saturation. In such cases, the early kinetics of *F*_*norm*_(*t*) are better described by a linear function. This behavior can be modeled by approximating the right-hand side of Eq. (15) using the first-order Taylor series expansion of the exponential function around zero, *e*^*x*^ ≈ 1 + *x*, which is valid for sufficiently small values of *x*. Under this approximation, *F*_*norm*_(*t*) can be expressed as:

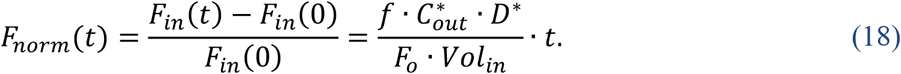

A very similar expression can be obtained by applying Taylor series expansion for background-subtracted recordings from tracer release assays, presented in Eq. (16):

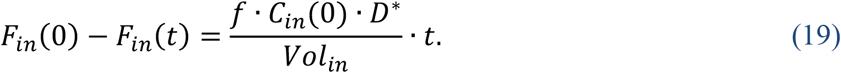

Thus, Eqs. (18)-(19) predict that the normalized and/or background-subtracted fluorescence intensity in tracer uptake and release assays initially follows a linear relationship of the form:

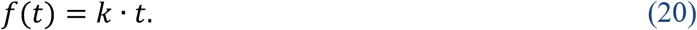

A representative example of such a case, obtained from DAPI uptake assay, is shown in Fig. 1.

**Fig. 1.**
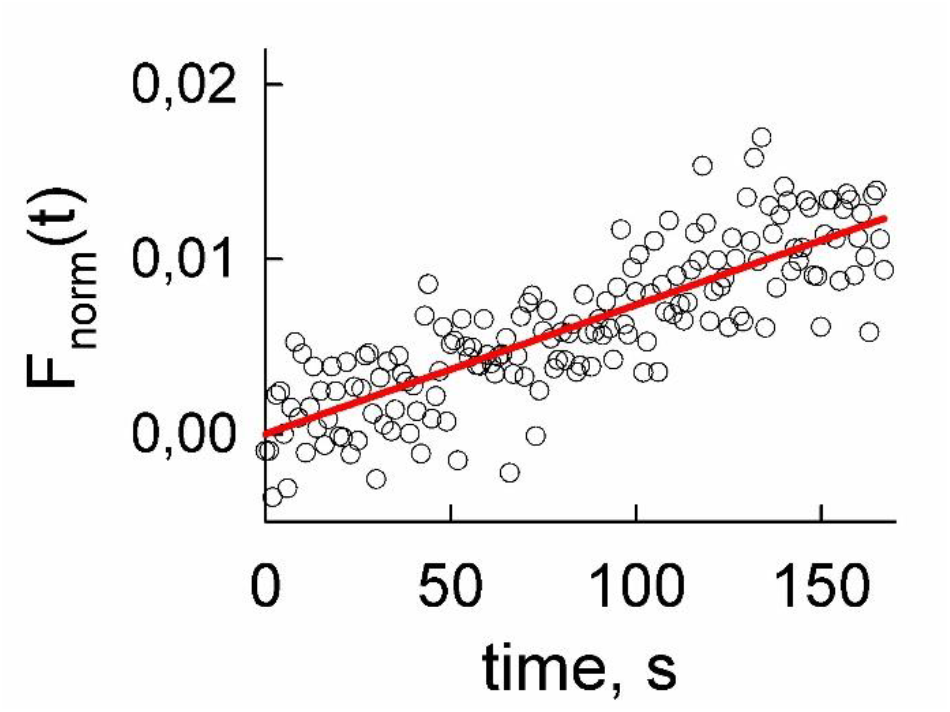
Representative example of changes in normalized intracellular fluorescence intensity, *F*_*norm*_(*t*), during a DAPI permeation assay. At the initial phase of the experiment, the recorded signal (open black circles) show no indication of saturation and is well-fitted by a linear function *F*_*norm*_(*t*) = *k* · *t* with *k* = 7.38·10^−5^ s^-1^.

In Eq. (20), the slope *k* reflects the quantity 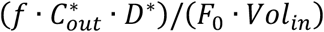 or 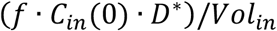, which has the dimension of a rate or a rate multiplied by fluorescence intensity, respectively. Importantly, the slope *k* can be directly compared to the product *a*·*b* from the exponential function *f*(*t*) = *a*·(1 – *exp*(*-b·t*)), as both *k* and *a*·*b* describe the same physical quantity. However, this comparison is valid only when the saturation of *F*_*norm*_(t) is determined primarily by diffusion. In certain cases, such as with DAPI, which fluoresces upon binding to intracellular nucleic acids, saturation may instead be limited by the availability of binding sites rather than diffusion. To avoid this confounding factor, our analysis focuses only on the initial linear segments of the data.

Thus, using a kinetic model to describe changes in fluorescence intensity in tracer uptake assays allows us to estimate the parameter 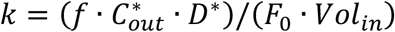. Using previous definitions for 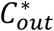 and *D* ^*^, as well as Eqs. (2)-(4), *k* can be expressed as follows:

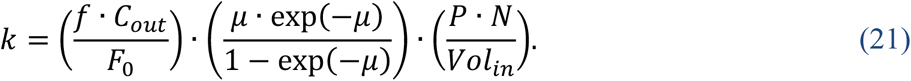

Here the initial fluorescence background level, *F*_0_, is directly obtained from imaging experiments, as is the concentration of tracer molecules in the bathing medium, *C*_*out*_. Parameter 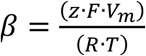 depends on resting membrane potential *V*_*m*_, which can be evaluated from independent electrophysiological recordings. Thus, we can define the estimable quantity *K*

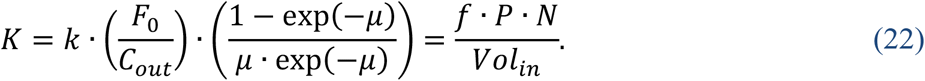

For tracer release assays, the equivalent expression for *K* can be obtained by removing *F*_0_ and replacing *C*_*out*_ with *C*_*in*_(0). The relative fluorescence intensity per concentration unit, *f*, in principle, cannot be easily determined during a tracer uptake experiment, as it depends not only on the fluorophore properties but also on specific experimental conditions. However, if the same conditions are maintained across different imaging experiments, *f* can be assumed constant, as it should not depend on the type of permeable channels. Consequently, the estimated quantity *K* should be proportional to the product of single-channel permeability *P* and total conductance *g*, and inversely proportional to cell volume *Vol*_*in*_.

### 2. Cell segmentation and fluorescence imaging data extraction using computer vision

Automated segmentation of HeLa cell boundaries was performed using an AI-based cell detection system (detailed in Methods section). It enabled automatic extraction of the changes in fluorescence intensity over time, as well as evaluation of cell surface area and estimation of cell volume for the dye uptake experiments.

First, the bright-field images were analyzed by the AI model to identify individual cell boundaries (see a representative example in Fig. 2A and B). Following the segmentation, cell boundaries from bright-field images were overlapped onto fluorescence dark-field images acquired using EGFP filter sets to visualize Cx26-msfGFP or Cx26-A49E-msfGFP expression. To evaluate Cx expression levels, we quantified the average pixel intensity within each automatically segmented cell region. Cells with mean fluorescence intensities above a defined threshold were considered as expressing Cx (successfully transfected) and were highlighted with green boundaries. In contrast, cells that did not exhibit significant fluorescence, were deemed as non-transfected and were highlighted with red boundaries (Fig. 2C).

**Fig. 2.**
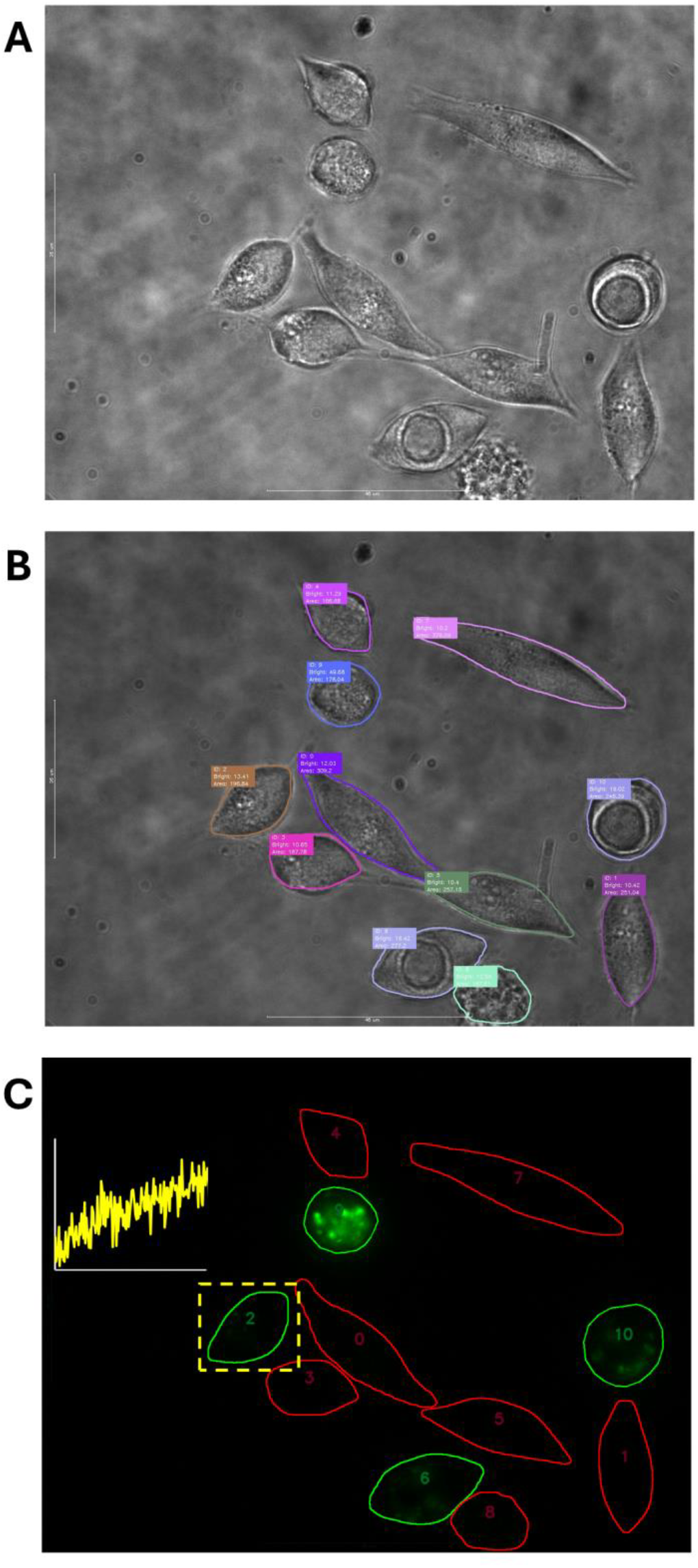
Automated cell segmentation and quantification of dye uptake in connexin-expressing HeLa cells. **(A)** A fragment of a representative bright-field image used to identify cell boundaries. **(B)** The same bright-field image as in (A), but with identified cell boundaries, denoted by solid lines of difference colours. Cell segmentation was performed an AI-based Mask R-CNN model. Additional cell parameters, enable by cell identification, such as measured surface area measurement are shown nearby individual cells. **(C)** Fluorescence dark-field image showing Cx expression, which was obtained using EGFP filter sets. Cells with mean fluorescence intensity above a set threshold, indicating Cx expression, are highlighted in green, while non-transfected cells are outlined in red. The yellow curve represents the time course of DAPI uptake within the marked cell region (yellow dashed rectangle).

Subsequent analysis focused on Cx-expressing cells that evaluated DAPI uptake by calculating the changes in the average pixel intensity within each defined cell boundary over time. Fig. 2C shows a representative example of fluorescence changes over time (solid yellow line in a graph) extracted from the Cx-expressing cell denoted by the dashed rectangular region.

### 3. The relationship between Cx hemichannel conductance and fluorescence intensity of EGFP inside segmented cells

First, we examine whether the fluorescence intensity of EGFP-tagged Cx protein could serve as a proxy for the number of functional Cx hemichannels. If successful, this approach could significantly reduce the need for time-consuming electrophysiological measurements, thereby streamlining data analysis.

To evaluate the relationship between fluorescence intensity and hemichannel conductance, we performed whole-cell patch-clamp recordings on single, isolated cells. To promote hemichannel opening and prevent their inhibition by extracellular divalent cations, cells were incubated for 5 minutes in a divalent cation-free solution, which was also used for continuous perfusion during the recording. Hemichannel currents were measured right at the onset of a voltage step, providing an estimate of the hemichannel conductance at the resting membrane potential, *g*. Dividing this overall conductance by the unitary conductance γ yielded an estimate of the number of functional Cx hemichannels, *g*/γ.

Simultaneously, we captured bright-field and dark-field images of each cell. Bright-field images were used to for AI-based cell boundary detection. The dark-field images were excited using EGFP filters, which allowed us to identify Cx protein expression inside the cell (see a representative example in Fig. 3A). We then computed the total EGFP fluorescence intensity (FI_EGFP_) by summing the pixel intensities within the segmented cell boundaries. This allowed us to examine the relationship between FI_EGFP_ and number of functional hemichannels.

**Fig 3.**
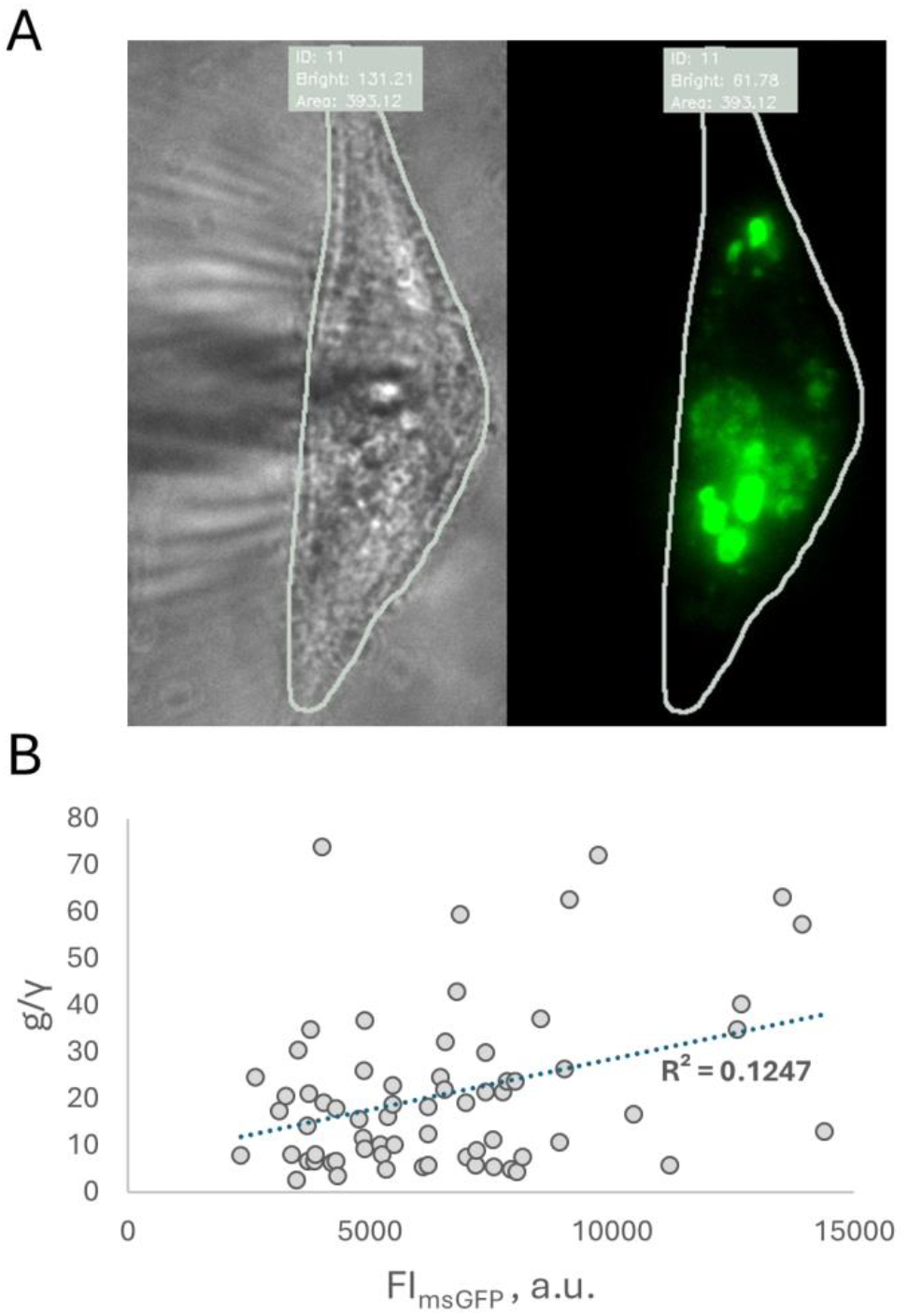
Number of functional Cx hemichannels cannot be reliably predicted by fluorescence intensity data. **(A)** Representative images of a cell expressing WT Cx26-EGFP protein. Left panel shows bright-field image displaying cellular morphology. Right panel shows dark-field image showing EGFP fluorescence indicating Cx26-EGFP protein expression. The cell boundary (green contour) was obtained using an AI-based segmentation model. **(B)** The relationship between fluorescence of EGFP protein, FI_EGFP_, the ratio of overall and unitary Cx hemichannel conductances, which reflects the number of functional Cx hemichannels in a cell. Dashed line represents a best-fit linear model together with the determination coefficient, R^2^.

Fig. 3B shows the g/γ - FI_EGFP_ relationship in cells expressing either WT Cx26 or the mutant Cx26^*^A49E. A visual inspection reveals a weak to moderate linear correlation between g/γ and FI_EGFP_, which was confirmed by a statistically significant but relatively low Pearson correlation coefficient, (ρ = 0.35, p-value = 0.0036). Most notably, the coefficient of determination for a fitted linear model was very low (R^2^ = 0.125), indicating that only ∼12.5 % of the variance in functional channel number can be explained by fluorescence intensity. This finding strongly suggests that FI_EGFP_ is not a suitable proxy for functional hemichannel number, and direct electrophysiological measurements remain necessary.

This result is consistent with the biological complexity of Cx hemichannel expression. Total cellular fluorescence reflects not only functional, membrane-inserted Cx hemichannels but also includes non-functional pools, such as misfolded subunits, proteins undergoing trafficking, and channels targeted for degradation. Additionally, even membrane-localized maybe non-conductive due to gating state or post-translational modifications. These factors collectively contribute to the observed disconnection between abundance of Cx protein and functional hemichannel conductance.

### 4. Evaluation of cell volume using data from cell segmentation and imaging system

To quantity fluorescence intensity changes during tracer uptake assays, we employed a kinetic ODE model in which the parameter *K* = (*f* · *P* · *N*)/*Vol*_*in*_, represents a composite rate constant, there *f* denotes fluorescent per unit of concentration, *P* is single-channel permeability, *N* is the number of open channels, and *Vol*_*in*_ is the intracellular volume. While some previous studies assumed that *Vol*_*in*_ remains constant [25] or is unaffected by the type of expressed channels [20], we sought to explicitly evaluate *Vol*_*in*_ and its influence on assessing Cx hemichannel permeability.

Accurate cell volume measurements typically require specialized imaging techniques such as 3D confocal microcopy. However, for adherent cells grown on glass coverslips, like those used in our DAPI uptake assay, *Vol*_*in*_ can be approximated based on the cell surface area *A*_*in*_ obtained from fluorescent imaging segmentation. These cells typically spread over the coverslip surface, forming a shape that can be approximated as a prism with an irregular base. Under this assumption, cell volume can be estimated as:

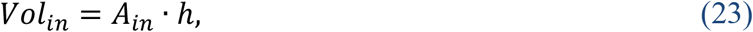

where *h* is the average cell height.

Although real cell morphology is complex, with height varying across the cell (e.g., HeLa cells can range from submicron heights near the periphery to ∼4.5 μm at the center [26]), using an average height provides a practical approximation. Therefore, even if local height varies, the total volume should scale proportionally with the measured surface area.

Using our cell segmentation data, we calculated surface areas and estimated *Vol*_*in*_, assuming an average cell height of ∼3 μm. To evaluate the relevance of these estimates for permeability analysis, we examined the relationship between *Vol*_*in*_ and the parameter *K*.

If *Vol*_*in*_ is estimated accurately and is independent of the product *f* · *P* · *N*, then *K* should be inversely proportional to *Vol*_*in*_, implying a hyperbolic relationship:

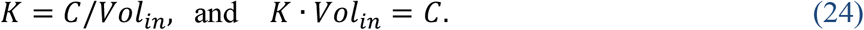

Conversely, if *Vol*_*in*_ estimates are inaccurate or strongly correlated with *f* · *P* · *N*, then *K* would appear constant in *K* vs *Vol*_*in*_ plot, and the product *K*·*Vol*_*in*_ would increase linearly with *Vol*_*in*_:

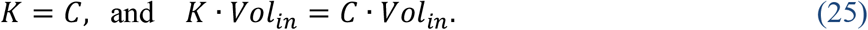

The upper and middle panels in Fig. 4 illustrate these two theoretical models using simulated data. Random variations in *N, Vol*_*in*_, and permeability constants *P* where adjusted to match the observed experimental variability. In Model 1 (upper panels), *N* and *Vol*_*in*_ were generated as independent variables. In Model 2 (middle panels), they were positively correlated (correlation coefficient 0.65), using Cholesky decomposition. In each case, the best-fit model (red lines) was selected using Akaike’s Information Criterion (AIC).

**Fig. 4.**
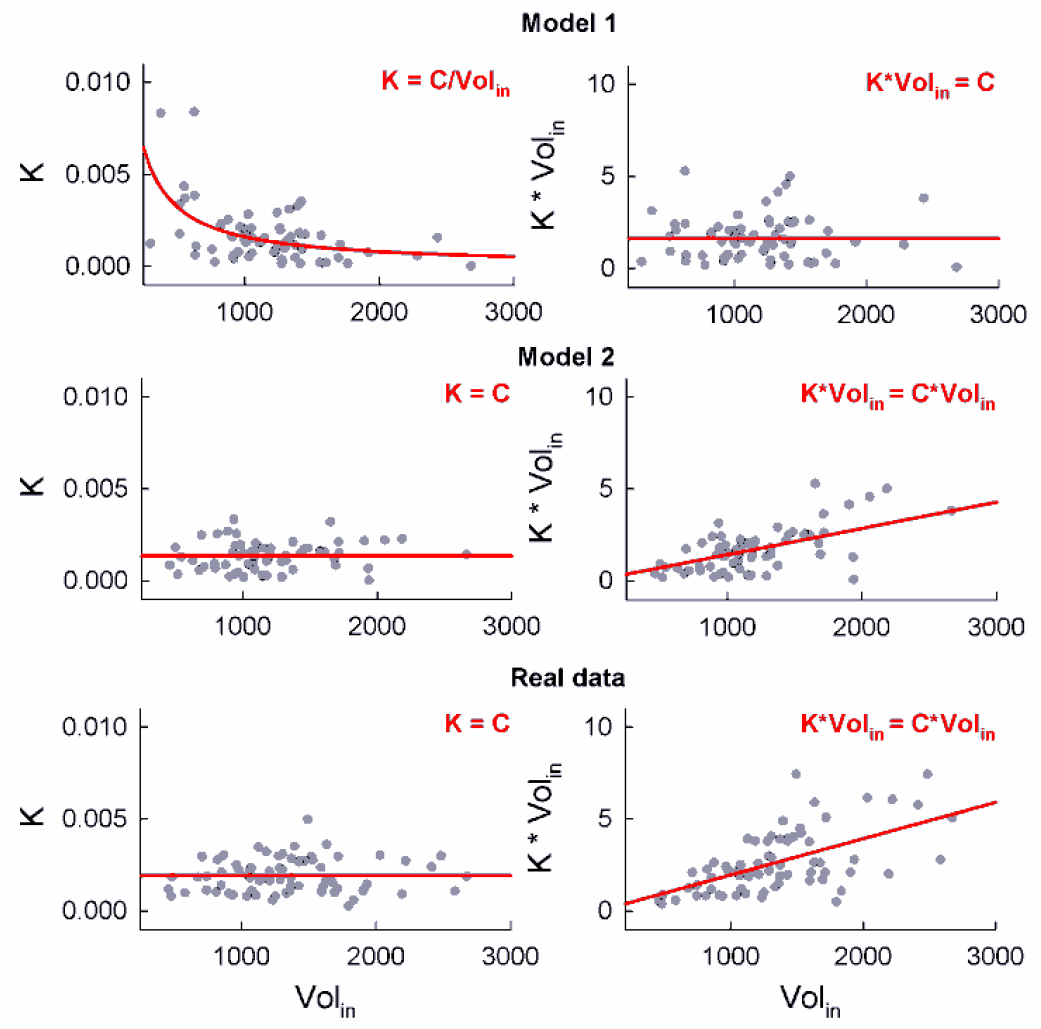
Relationship between estimated slope parameters *K* and cell volumes *Vol*_*in*_. The upper panels show fits of two alternative models (solid red lines) to simulated datasets (grey circles): *K* versus *Vol*_*in*_ (left) and *K*·*Vol*_*in*_ versus *Vol*_*in*_ (right). The lower panel shows presents model fits to experimental data. The results support the second model, suggesting that estimated *K* does not depend on surface area-based estimates of *Vol*_*in*_. Units: *Vol*_*in*_ in μm^3^, *K* in a.u.· L/(μM·s).

The lower panel of Fig. 4 presents fits to experimental data. The linear model (see Eqs. (18) and (20)) was applied to normalized fluorescence traces from HeLa cells transfected with Cx26, and *K* was estimated using Eq. (22). The average resting membrane potential in transfected cells was near zero with small standard deviation (*V*_*m*_ = -8.22 ± 1.77 mV, sample size 15). According to the GHK model prediction, this negative membrane potential should enhance DAPI uptake for ∼35 percent compared to for ∼35 percent compared to *V*_*m*_ = 0 mV resting potential, given DAPI’s valence *z* = +2.

Experimental data favored the constant *K = C* model in the *K* vs. *Vol*_*in*_ plot (AIC: = -587.63 and -634.55 for Model 1 and 2, respectively), while the linear model *K*·*Vol*_*in*_ = *C*·*Vol*_*in*_ best fit the *k*·*Vol*_*in*_ vs. *Vol*_*in*_ plot (AIC: 416.15 and 385.87 for Model 1 and 2, respectively). These findings support Model 2, indicating that *K* is independent of our estimates of *Vol*_*in*_. Similar results were obtained from HeLa cells transfected with the Cx26^*^A49E.

We propose two possible explanations for this result, each with different implications. First, the surface area-based volume estimates derived from AI-assisted segmentation may lack accuracy. Cell with greater spread may have reduced height, weakening the correlation between surface area and actual volume. Representative examples of atomic force microscopy reconstructions and height measurements of HeLa cells at different phases of cell cycle presented in [26] provides some support for this hypothesis. If true, then *Vol*_*in*_ estimates may be unreliable and should be excluded from the further analysis.

Alternatively, even if volume estimates are accurate, *Vol*_*in*_ may be positively correlated with the number of open channels, *N*. For instance, if channel density is uniform across the membrane, then the larger surface area would imply a higher *N*. Our simulations indicate that a Volin-N correlation of ¬0.5 can explain the observed model fit differences presented in Fig. 4. In our own data this correlation was a little lower (∼0.36) but statistically significant. Thus, if the deviation from the inverse proportionality between *K* and *Vol*_*in*_ arises from this correlation, incorporating *Vol*_*in*_ into further analysis may still be informative. We address this possibility using our proposed statistical model in the following section.

### 5. Statistical test for comparing permeabilities of different channels using fluorescent imaging and electrophysiological data

Fluorescence imaging data provides an estimable quantity *K*, which is proportional to the product of single-channel permeability *P* and total conductance *g*. If the total conductance *g* is not measured simultaneously with fluorescence imaging data, direct comparison of estimated values of *K* does not necessarily reveal differences in *P*, as variations in *K* may also reflect differences in the number of open channels.

For example, consider a comparison between wild-type (WT) and mutant channels. A mutation may affect not only single-channel permeability, but also channel expression levels and/or channel trafficking, which would influence the number of functional channels *n*. Additionally, mutant channels may exhibit altered open probability *p*_*o*_, leading to changes in the number of open channels *N* = *g*/*γ*.

Thus, for a direct comparison of permeability, it is necessary to separate the effects of single-channel permeability from the number of available channels. We propose that this can be achieved through statistical modeling.

First, we assume that single channel permeability, *P*, remains constant for channels of the same type across different experiments. This assumption of identical independent channels is commonly used in mathematical models of ion channel behavior. However, the estimated number of open channels, *N*, can vary significantly between experiments and should be treated as a random variable.

To determine differences in channel permeability *P* from estimates of the quantity *K*, we propose incorporating independent electrophysiological measurements of *N*. These measurements should be obtained under conditions matching those used in the fluorescent imaging experiments. For instance, when evaluating tracer uptake through open connexin hemichannels, electrophysiological recordings should be conducted under the same extracellular divalent cation concentration. Additionally, the same cell lines should be used to minimize variability in the number of functional channels.

For direct comparison of WT and mutant channels, with permeabilities *P*_*wt*_ and *P*_*mut*_, respectively, we incorporate independent datasets for functional hemichannel measurements (*N*_*wt*_ and *N*_*mut*_) along with the estimated *K* values (*K*_*wt*_ and *K*_*mut*_).

### Challenges in direct comparison

#### 1) Assumption of identical conductance distributions

If *N*_*wt*_ and *N*_*mut*_ follows the same distribution, direct comparison of *K*_*wt*_ and *K*_*mut*_ would be valid, as any observed differences would reflect differences in *P*_*wt*_ and *P*_*mut*_. However, this approach requires two consecutive statistical tests– first, to compare the distribution of *N*_*wt*_ and *N*_*mut*_, and second, to compare *K*_*wt*_ and *K*_*mut*_. This increases the probability of false positives or negatives. Thus, the p-values in these two steps should be adjusted accordingly, e.g., for an overall significance level of 0.05, each individual test should use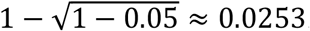.

#### 2) Differences in conductance distributions

If *N*_*wt*_ and *N*_*mut*_ differ significantly (e.g., due to altered open state probability, expression levels or trafficking of mutant channels), direct comparison of *K*_*wt*_ and *K*_*mut*_ may be misleading. If *N*_*wt*_ is systematically higher that *N*_*mut*_, higher mean values of *K*_*wt*_ does not necessarily imply that *P*_*wt*_ is greater than *P*_*mut*_.

To address these limitations, we propose a statistical model based on likelihood ratio test.

### Likelihood ratio test for channel permeability

The likelihood ratio test evaluates whether a more complex model provides a significantly better fit to the data compared to a simpler model, which is a special case (submodel) of the expanded model. The submodel has fewer parameters and necessarily provides a lower likelihood estimate. However, if the likelihood gain from the expanded model is insignificant, the simpler model is preferred. This can is quantified using the ratio of likelihoods:

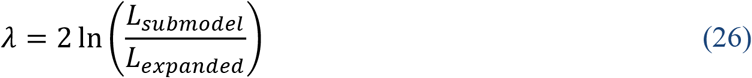

Under the null hypothesis (i.e., the submodel fits data as well as the expanded model), λ follows a chi-squared distribution with degrees of freedom equal to the difference in parameter count between the models.

### Formulation of the model

Since *K*_*wt*_ is proportional to *N*_*wt*_, we express it as *K*_*wt*_ = *C*_*wt*_ · *N*_*wt*_, where *C*_*wt*_ is assumed to be a constant. Similarly, for mutant channels *K*_*mut*_ = *C*_*mut*_ · *N*_*mut*_.

If *N*_*wt*_ follows a probability distribution with a cumulative distribution function *F*_*wt*_(*x; θ* _*wt*,1_, … *θ* _*wt,m*_), then *K*_*wt*_ follows a probability distribution with a cumulative distribution function *F*_*wt*_ (*x*⁄ *C*_*mut*_ ; *θ* _*wt*,1_, … *θ* _*wt,m*_). Similarly, for *N*_*mut*_ and *K*_*mut*_ the respective cumulative distribution functions are given by *F*_*mut*_ (*x; θ* _*wt*,1_, … *θ* _*wt,n*_) and *F*_*mut*_ (*x*⁄ *C*_*mut*_ ; *θ* _*wt*,1_, … *θ* _*wt,n*_). If *N*_*wt*_ and *N*_*mut*_ follows the same type of distribution, they would have the same number of parameters (e.g., *m* = *n*), although their values may differ.

Many common distributions, such as the gamma distribution, are stable under multiplication by a constant, preserving the functional form of *K*. For example, if *X* ∼*G* (*α, σ*) (gamma-distributed with shape *α* and scale *σ*), then multiplying *X* by a positive constant *C* results in another gamma-distributed random variable with the same shape parameter *α*, and scale parameter *C*·*σ*: *Y*∼*G*(*α C*·*σ*). Thus, if *N*_*wt*_ ∼*G*(*α*_*wt*_, *σ*_*wt*_), then N_*wt*_ ∼*G*(*α*_*wt*_, *C*_*wt*_ ·*σ*_*wt*_), preserving the distribution type.

### Maximum likelihood estimation

We define the submodel under the null hypothesis that assumes equal permeabilities:

- *N*_*wt*_ ∼*F*_*wt*_(*x; θ* _*wt*,1_, … *θ* _*wt,m*_);
- *K*_*wt*_ ∼*F*_*wt*_ (*x*⁄ *C*_*mut*_ ; *θ* _*wt*,1_, … *θ* _*wt,m*_);
- *N*_*mut*_ ∼*F*_*mut*_(*x; θ* _*mut*,1_, … *θ* _*mut,m*_);
- *N*_*mut*_ ∼*F*_*mut*_(*x*⁄ *C*_*mut*_ ; *θ*_*mut*,1_, … *θ* _*mut,m*_);
- *C*_*wt*_*=C*_*mut*_ = *C* (assuming equal permeability)

The model has 2*m*+1 parameters, assuming the same type of distributions for *g*_*wt*_ and *g*_*mut*_. In the expanded model, the assumption *C*_*wt*_*=C*_*mut*_ = *C* is removed, allowing for different permeabilities between WT and mutant channels. Thus, the expanded model has 2*m*+2 parameters.

After estimating likelihood *L*_*submodel*_ and *L*_*expanded*_, we compute 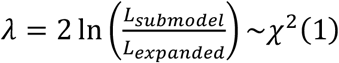. Under the null hypothesis, p-value is given by: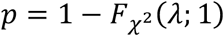. If *p* < 0.05, we reject the null hypothesis, indicating significant difference in permeabilities.

### Model validation

To ensure the validity of the model, the estimated parameters should fit empirical data well. This can be assessed using standard nonparametric tests, such as the Kolmogorov-Smirnov test, to compare the observed data distributions with those predicted by the model.

### Evaluation of cell volume data

We propose that the statistical model can also serve as a tool to assess the quality of surface-area based estimates of cell volume, *Vol*_*in*_. If these estimates reflect true cell volumes, and the deviation from the expected relationship *K* = *C*/*Vol*_*in*_ (with constant *C*) arises primarily from a correlation between *Vol*_*in*_ and a number of open channels *N*, then incorporating *Vol*_*in*_ into the model – specifically, by replacing the *K* data sets with *K*·*Vol*_*in*_ – should improve the fit. In our framework, the composite rate constant *K*=(*f* · *P* · *N*) */Vol*_*in*_ is assumed to be proportional to *N*, other factors being constant. However, *Vol*_*in*_ varies across individual traces, even within the same cell line. If not accounted for, this variation may introduce additional noise, leading to discrepancies from the assumed relationship *K* = *C* · *N*, which in turn constrains the joint distribution of *N* and *K* (for example, to the same distribution family such as gamma). Therefore, if incorporating *Vol*_*in*_ into the model improves goodness-of-fit (for example, by increasing p-values in Kolmogorov-Smirnov tests or likelihood values), this would support the validity and informativeness of the obtained *Vol*_*in*_ estimates.

### 6. Application of the proposed methodology to experimental data

We applied the proposed methodology to electrophysiological and fluorescence imaging data obtained from DAPI uptake assays in hemichannels formed by WT Cx26 and the Cx26^*^A49E variant.

First, we measured the total conductance of WT Cx26 and Cx26*A49E hemichannels using whole-cell patch-clamp recordings. These measurements were performed near 0 mV membrane voltage in divalent cation-free solutions to replicate the conditions used in DAPI uptake assays. A unitary conductances of ∼340 pS was used, as reported in [27]. Dividing the measured conductance by unitary conductance, g/γ, gave us the estimates of open hemichannels, *N*. In total, we obtained 40 measurements of *N* for WT Cx26 and 40 for Cx26^*^A49E hemichannels.

In DAPI uptake assays, we recorded time courses of fluorescence intensity. Each background-subtracted and normalized time course was fitted using the proposed kinetic model. Linear fits described in Eqs. (18) and (20) yielded the rate parameter *K* (see Eq. (22)) for each time course. In total, we obtained 70 estimates of *K* for WT Cx26 and 36 for Cx26^*^A49E, hemichannels.

To analyze the datasets *N*_*Cx26*_, *g*_*Cx26*A49E*_, *K*_Cx26_, and *K*_*Cx26*A49E*_, we applied the proposed statistical model based on the likelihood ratio test. A gamma distribution was chosen, as it provided a good fit for all four datasets when fitted independently from each other. In the restricted submodel, the permeability constants of Cx26 and Cx26*A49E, denoted as *C*_Cx26_ and *C*_Cx26*A49E_, were assumed to be equal, leading to the following five-parameter submodel:

- *N*_*Cx*26_ ∼*G*(*x*; *α*_*Cx*26,_ *σ*_*Cx*26_);
- *K*_*Cx*26_ ∼*G*(*x* /*C*_*Cx*26_; *α*_*Cx*26,_ *σ*_*Cx*26_);
- *N*_*Cx*26 * *A*49*E*_ ∼*G*(*x*; *α*_*Cx*26 * *A*49*E*,_ *σ*_*Cx*26 * *A*49*E*_);
- *N*_*Cx*26 * *A*49*E*_ ∼*G*(*x* /*C*_*Cx*26* *A*49*E*_ *α*_*Cx*26 * *A*49*E*_ *σ*_*Cx*26 * *A*49*E*_);
- *C*_*Cx*26_= *C*_*Cx*26* *A*49*E*_=*C*.

In contrast, the expanded model allowed for permeability coefficients to differ from each other. Thus, eliminating the condition *C*_*Cx*26_= *C*_*Cx*26* *A*49*E*_=*C* resulted in a six-parameter statistical model. These models were fitted to the data using the maximum likelihood criterion.

To test the validity of the surface-area-based cell volume estimates, we also evaluated volume-adjusted permeability values, *K*·*Vol*_*in*_. For the Cx26^*^A49E hemichannels, two outliers were excluded from the obtained *K*·*Vol*_*in*_ data set using a conservative criterion of six median absolute deviations from the median. Estimated parameters and corresponding log-likelihood (LL) are shown in Table 1.

**Table 1.** Estimated parameters of statistical models.

| | *N* and *K* | | *N* and $K \cdot Vol_{in}$ | |
| --- | --- | --- | --- | --- |
| | $C_{Cx26} = C_{Cx26*A49E}$ | $C_{Cx26} \neq C_{Cx26*A49E}$ | $C_{Cx26} = C_{Cx26*A49E}$ | $C_{Cx26} \neq C_{Cx26*A49E}$ |
| $\alpha_{Cx26}$ | 2.8806 | 3.1398 | 2.1386 | 2.4048 |
| $\sigma_{Cx26}$ | 5.8297 | 6.6420 | 7.3229 | 8.6720 |
| $C_{Cx26}$ | 0.000133 | 0.000093 | 0.2069 | 0.1259 |
| $\alpha_{Cx26*A49E}$ | 1.0103 | 2.3829 | 1.3703 | 2.3104 |
| $\sigma_{Cx26*A49E}$ | 55.5155 | 7.3041 | 26.3297 | 7.5335 |
| $C_{Cx26*A49E}$ | 0.000133 | 0.000759 | 0.2069 | 0.6899 |
| *LL* | 169.39 | 213.91 | -571.08 | -540.9327 |

Likelihood ratio tests yielded a statistic λ = 89.04 using the *N* and *K* datasets, and λ = 60.30 when using *N* and volume-adjusted *K*·*Vol*_*in*_. In both cases, p-values were close to 0, indicating that the expanded model, which allows distinct permeabilities, provides a significantly better fit than the restricted model.

Further validation was performed using Kolmogorov-Smirnov test. The p-values obtained for the gamma distribution fits are listed in Table 2.

**Table 2.** Kolmogorov-Smirnov test p-values.

| Dataset | $C_{Cx26} = C_{Cx26*A49E}$ | $C_{Cx26} \neq C_{Cx26*A49E}$ |
| --- | --- | --- |
| $N_{Cx26}$ | 0.3498 | 0.1811 |
| $K_{Cx26}$ | 0.0880 | 0.7234 |
| $N_{Cx26*A49E}$ | $5.26 \cdot 10^{-6}$ | 0.6905 |
| $K_{Cx26*A49E}$ | $2.53 \cdot 10^{-7}$ | 0.2326 |
| $N_{Cx26}$ | 0.1049 | 0.4968 |
| $K_{Cx26} \cdot Vol_{in}$ | 0.0014 | 0.8495 |
| $N_{Cx26*A49E}$ | 0.0028 | 0.7130 |
| $K_{Cx26*A49E} \cdot Vol_{in}$ | $4.20 \cdot 10^{-12}$ | 0.6336 |

The Kolmogorov-Smirnov test results clearly show that the expanded model, which allows different permeability constants, provides substantially better fits. For the expanded model, all p-values exceeded 0.05, indicating adequate agreement with the data. In contrast, the restricted model exceeded this threshold only for the unadjusted *K* values in the Cx26 datasets and for *N*_*Cx26*_ when combined with volume-adjusted *K*·*Vol*_*in*_.

Furthermore, volume-adjusted *K*·*Vol*_*in*_ values yielded higher p-values across datasets, indicating better model fits. They also produced lower AIC values for both *N*_*Cx26*_ and *N*_*Cx26*A49E*_ datasets (*K* and *K*·*Vol*_*in*_ datasets do not allow for direct comparison due to different scales). For example, using unadjusted *K*, AIC values were 499.46 and 368.91, whereas with volume-adjusted data, they were only 394.35 and 359.26 (differences greater than 2 units are considered statistically significant, lower values indicating better fit). Thus, the improved fits suggest that the volume-adjusted variable *K*·*Vol*_*in*_ more accurately preserves the expected distribution, as predicted by the relationships *K*·*Vol*_*in*_ = *C*·*N*, compared to the unadjusted *K* values predicted by *K* = *C*·*N*. This supports the hypothesis that surface-area-based *Vol*_*in*_ estimates are informative proxies for actual cell volumes.

The theoretical cumulative distribution function fits are presented in Fig. 5.

**Fig. 5.**
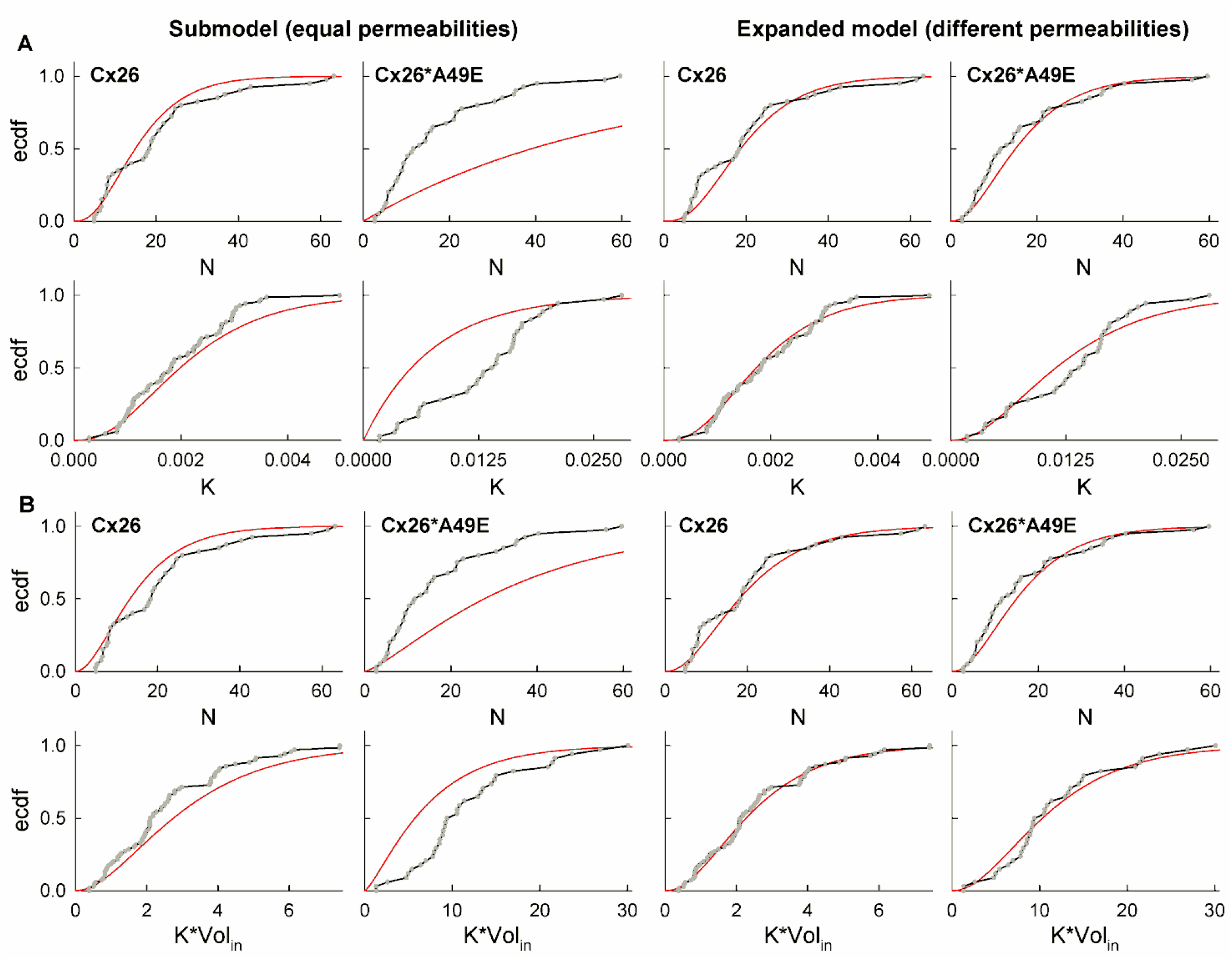
Fits of gamma distribution using the proposed statistical model. **(A)** Fits to Cx26 and Cx26*A49E datasets using the restricted model (left panels) and expanded model (right panes). Grey circles and black lines represent empirical cumulative distribution functions (ecdfs); red lines indicate fitted gamma cumulative distribution functions. (B) Corresponding fits for datasets using permeability rate *K* values adjusted by estimated cell volumes *Vol*_*in*_. Units of *K* are in a.u.· L/(μM·s), and Volin was measured in μm^3^.

Overall, these results demonstrate that WT Cx26 and Cx26^*^A49E hemichannels exhibit significantly different permeabilities to DAPI. The expanded model predicts that Cx26^*^A49E hemichannels are more permeable, showing more than an eightfold increase using unadjusted K, and more that a fivefold increase using volume-adjusted *K*·*Vol*_*in*_. This finding is consistent with previous studies indicating that the negatively charged ring near position 49 is critical for permeability in Cx26 and Cx30 hemichannels [22; 27]. The neutral-to-negative substitution at this site enhances Cx26 permeability to positively charged molecules such as DAPI.

### 7. Statistical power of the proposed statistical model to detect differences in permeability

To evaluate the sensitivity of the proposed methodology in detecting differences in permeability, we performed a simulation-based power analysis of the statistical test. In these simulations, we generated sets of random variables *X* and *Y*, correspond to the conductances of different channel types of channels, such as WT and mutant channels (*N*_*w*t_ and *N*_*mut*_). The associated permeability datasets, *PX* and *PY*, where generated using the relationship *PX* = *P*_*X*_ *· X* and *PY* = *P*_*Y*_ *· Y*, where *P*_*X*_ and *P*_*Y*_ are positive constants representing permeability coefficients of different channels.

For simplicity, we modeled *X* and *Y* using gamma distribution with identical equal shape and scale parameters (α_X_ = α_Y_ and *σ*_*X*_ = *σ*_*Y*_), selected to reflect values observed in experimental data. The permeability coefficient *P*_*X*_ was fixed to 1, while *P*_*Y*_ was varied to obtain different ratios *P*_*Y*_/*P*_*X*_. According to the properties of random variables gamma, permeability datasets *PX* and *PY* were described by a shape parameter α_PX_ = α_PY_ = α_X_ = α_Y_ and scale parameters *σ*_*PX*_ = *P*_*X*_ · *PX* and *σ*_*PY*_ = *P*_*Y*_ · *PY*. This allowed us to assess the probability of detecting a statistically significant difference between *P*_*X*_ and *P*_*Y*_ using the likelihood ratio test. For each sample size, we generated 1000 independent datasets of *X, Y, PX*, and *PY* and applied the likelihood ratio test. The fraction of cases where the test yielded p-values below 0.05 was taken as the statistical power (probability 1-*β*, where *β* denotes statistical error of type II).

Simulation results in Fig. 6A show the dependence of test power on sample sizes *n*_*X*_ and *n*_*Y*_ (left panel) and *n*_*PX*_ and *n*_*PY*_ (right panel). In Fig. 6, the gamma distribution shape parameters were set to α_X_ = α_Y_ = 2.4, and the scale parameters to *σ*_*X*_ = *σ*_*Y*_ = 8.0. The power of the test increased similarly with increases in either *n*_*X*_ and *n*_*Y*_, or *n*_*PX*_ and *n*_*PY*_. Since *n*_*PX*_ and *n*_*PY*_ correspond to number of fluorescence intensity recordings in our methodology – data that is easier to collect in large amounts compared to electrophysiological conductance measurements – increasing *n*_*PX*_ and *n*_*PY*_ appears to be a more practical strategy for improving test power.

**Fig. 6.**
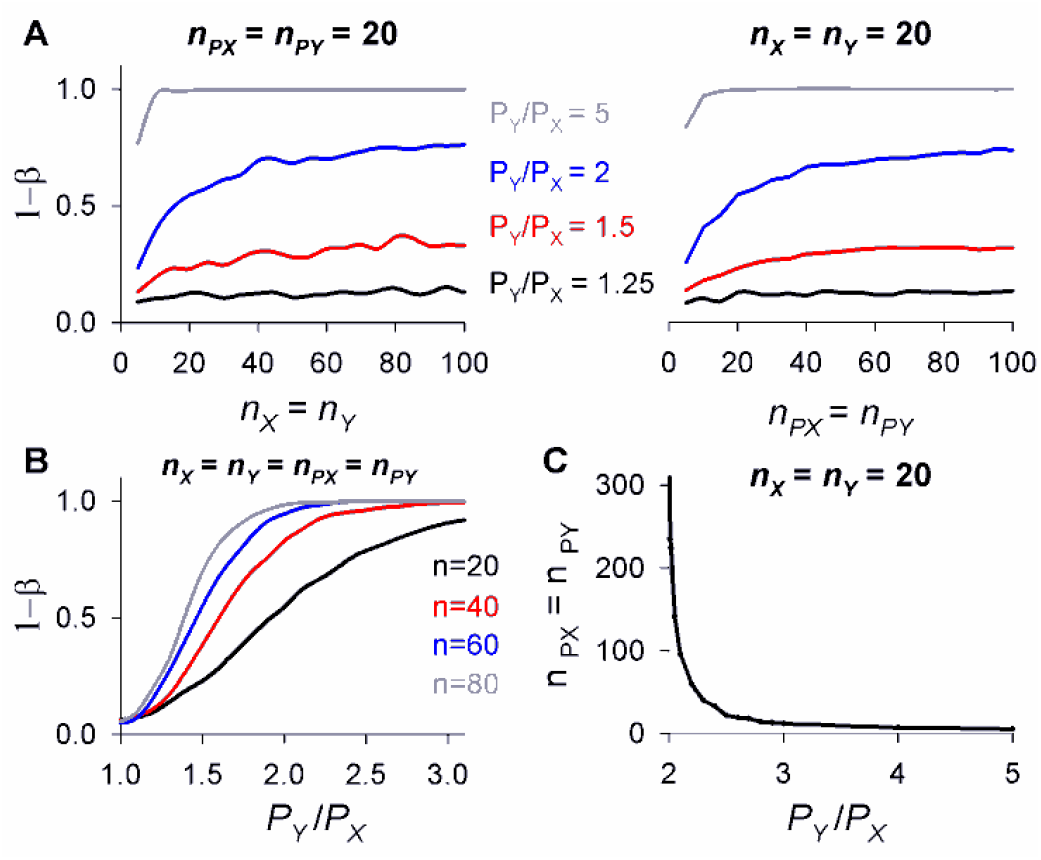
Evaluation of the statistical power of the proposed method for comparing Cx hemichannel permeability. **(A) Test power as a function of sample sizes of** hemichannel conductance estimates (*n*_*X*_ and *n*_*Y*_) and fluorescence intensity traces (*n*_*PX*_ and *n*_*PY*_) at varying permeability ratios *P*_*Y*_/*P*_*X*_. (**B**) Test power curves as a function of the permeability ratio *P*_*Y*_/*P*_*X*_, evaluated for different sample sizes. (**C**) Required sample sizes of fluorescence intensity traces to achieve a statistical power of 0.80 across different *P*_*Y*_/*P*_*X*_ ratios.

Fig. 6B illustrates the relationship between test power and the permeability ratio P_Y_/P_X_ at fixed sample sizes. These data suggest that while the proposed test cannot reliably detect small permeability differences, it achieves acceptable power (∼0.80 probability) for moderate sample sizes then ratio *P*_*Y*_/*P*_*X*_ > 2. Fig. 6C further confirms this by showing the minimum required sample size of *P*_*X*_ and *P*_*Y*_ (with *n*_*X*_ = *n*_*Y*_ = 20) needed to achieve a test power of 0.80. The required sample size decreases sharply once *P*_*Y*_/*P*_*X*_ exceeds 2. For larger permeability differences, such as those observed in our study (where estimated *P*_*Y*_/*P*_*X*_ > 5), the required sample size is relatively small (around 5-7 samples).

Our simulation data also demonstrated that these findings hold across different underlying distributions of *X* and *Y* (for example, varying shape α or scale β resulted in similar trends). Thus, this power analysis confirms that the proposed statistical test is robust in detecting permeability differences in electrophysiological and fluorescence imaging data. In our case, the observed permeability difference between WT Cx26 and Cx26^*^A49E hemichannels is large to be reliably detected by the proposed statistical approach.

## Discussion

We propose a methodology that enables the comparing of molecular permeability among different types of Cx hemichannels. The key feature of our approach is the integration of independently collected electrophysiological and fluorescence imaging data. Electrophysiological recordings provide estimates of the ratio of overall channel conductance to unitary conductance *g/γ*, serving as a proxy for the number of open channels, *N*. Fluorescence intensity kinetics yields estimates of the product *P*·*N*, where *P* represents the permeability of a single channel. By integrating data from different hemichannel types into a unified statistical model based on likelihood ratio test, our methodology allows for rigorous comparisons of permeability differences.

### Advantages, limitations, and potential improvements

#### Separating electrophysiological and fluorescent imaging data

The proposed methodology is technically less demanding than previously published techniques that simultaneously measure the number of channels and diffusion rates within the same cells [16; 17; 28]. Performing electrophysiological and fluorescent imaging experiments separately increases experimental flexibility, allowing these measurements to be conducted at different times and locations. As was noted in [29], while tracer uptake and release assays can be performed using standard equipment found in a typical molecular and cell biology lab, electrophysiological recordings require more specialized expertise and instrumentation. Our approach, which enables the pooling of data from different laboratories or previously published studies, is valuable in this context.

To ensure the validity of pooled data, it is essential that the distribution of functional channel numbers in fluorescence imaging experiments remains consistent with that in electrophysiological recordings. This requires using the same cell lines for a given hemichannel type both experimental approaches, as differences in protein expression patterns and cell sizes can affect the number of functional channels. Additionally, experimental conditions influencing open-state probability – such as membrane voltage and extracellular divalent ion concentration – must be standardized. For Cx hemichannels in particular, the same extracellular Ca^2+^ and Mg^2+^ concentrations should be used in both electrophysiological recordings and tracer uptake/release assays.

Importantly, our statistical model does not require identical channel number distributions between different Cx hemichannel types. This flexibility allows for cross-comparisons between datasets obtained from different cell types. However, for each hemichannel type, fluorescence imaging and electrophysiological data must be derived from the same cell line. Additional considerations include differences in resting membrane potential, and potential differences in cell volumes.

#### Required sample sizes

A limitation of our methodology is that quite large datasets are needed to detect relatively small permeability differences. Simulations suggest that resolving permeability differences smaller than twofold requires datasets exceeding 500-1000 samples to achieve a statistical power of 0.80. In contrast, direct measurements of single-channel permeability via simultaneous electrophysiological and diffusion rate recordings likely require far fewer samples for comparable statistical significance.

However, this drawback can be partially offset by the technical ease of our methodology, particularly when permeability differences exceed twofold. Simulation data indicate that the statistical power of our test depend equally on the sizes of both electrophysiological and imaging datasets. Large fluorescence imaging datasets can be obtained efficiently using high-throughput tracer uptake/release assays, which are technically simple than electrophysiological recordings and allow the collection of 5-20 samples per experiment. Automated cell detection and segmentation software can further streamline data analysis. When permeability differences exceed threefold, our approach may enable faster collection of sufficient amount of data compared to more complex simultaneous measurement techniques.

#### Simplified diffusion model

Our methodology employs a simplified diffusion model based on Fick’s law and GHK current equation to describe Cx hemichannel permeability. A similar approach was used in previous studies on Cx hemichannel permeability [29]. However, this model does not account for certain mechanistic features that are relevant for tracer uptake/release assays.

First, it assumes that fluorescence intensity is directly proportional to intracellular tracer concentration, despite the fact that some tracers, such as DAPI, fluoresce upon DNA binding. As a result, the model is not suitable for describing fluorescence recordings that exhibit saturation, as it cannot distinguish between saturation limited by diffusion and that limited by binding. In such cases, a more advanced tracer uptake models incorporating intracellular binding kinetics and cellular morphology could be applied instead, as was proposed in [30].

To minimize the impact of binding-related discrepancies, we restricted our analysis to the short initial segments of the fluorescence time courses, which can be adequately described by the linear model presented in Eqs. (18) and (20). A simple analysis indicates that under these conditions, the proposed methodology remains valid.

To account for tracer binding to DNA or other intracellular substrate, denoted as *S*, we consider the concentration of the tracer-substrate complex, [*C*_*in*_-*S*]. During the initial stage of the experiment, when the total biding site concentration is much greater than the amount of bound tracer, and binding does not significantly deplete the intracellular free tracer concentration *C*_*in*_(*t*), the kinetics of [*C*_*in*_-*S*] complex concentration can be approximated by the following ODE:

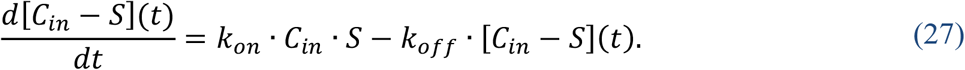

Here, *k*_*on*_ and *k*_*off*_ are the association and dissociation rate constants, respectively, and *S* is the concentration of available binding sites. Assuming that the initial concentration of the complex is zero, the solution of Eq. (27) is:

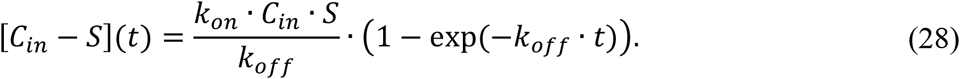

At early time points, when the product *k*_*off*_ · *t* is sufficiently small, the exponential term in the Eq. (28) can be approximated using a first-order Taylor series expansion, yielding the following linear function:

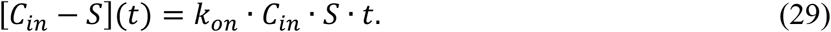

This result supports the use of our simplified linear model to describe the initial uptake phase, even when tracer binding occurs. Nevertheless, the inclusion of binding kinetics introduces additional parameters, such as *k*_*on*_ and *S*, which may contribute to variability in he estimated permeability rates. This should be taken into account when comparing hemichannel permeability across different experimental conditions or different cell lines.

#### Can fluorescent imaging data replace electrophysiological measurements of channel number?

Preliminary data suggest that fluorescence imaging, while correlated with Cx hemichannel currents, cannot yet serve as a direct substitute for electrophysiological recordings. This is expected, as total fluorescence emitted by Cxs tagged with fluorescent protein reflects the presence of channel proteins in the cell but does not necessarily correspond to functional channels inserted into the membrane, at least in 2D imaging data.

However, advances in machine learning and computer vision may enable future improvements. With sufficient training data linking fluorescence imaging to electrophysiological conductance measurements, artificial neural networks could be developed to estimate channel conductance form imaging data alone. This could be validated by kinetic modeling of diffusion rates. For example, comparing estimated diffusion rates against conductance values, similarly as we show for cell volume data in Fig. 3, could help assess the accuracy of such predictions.

#### Applicability to other types of channels

Our methodology, applied here to compare permeability of WT Cx26 and Cx26^*^A49E hemichannels, could also be extended to other large-pore channels permeable to small molecules. These include pannexin (Panx) channels, calcium homeostasis modulators (CALHMs), and volume-regulated anion channels (VRACs).

Similar to Cxs, Panxs form hemichannels permeable to small molecules, such as ATP [31]. Only three Panx isoforms have been identified in humans [32], but at least one is expressed in most mammalian tissues. Changes in Panx expression are associated with various pathological conditions in neural system [33; 34] and skeletal disorders [35; 36], and mutations in Panx1 genes were recently linked to genetic diseases, such as human oocyte death [37].

CALHM channels share functional properties with Cx hemichannels, such as voltage and Ca^2+^ sensitivity and permeability to small molecules [38; 39]. CALHM channels are important in neuronal excitability and muscle cell function [40]. Mutations in CALHM1 are associated with Alzheimer’s disease [41; 42], while CALHM3 channels play a role in taste perception [43; 44]. Interestingly, CALHM proteins can form oligomeric assemblies with varying protomer number [45; 46], each conceivably exhibiting differences in permeability.

VRAC channels, formed of leucine-rich repeat-containing (LLRC) proteins, regulate cell volume through Cl^-^ and osmolyte transport [47; 48; 49]. Mutations LRRC8A are linked to immune deficiency disorder agammaglobulinemia [50],[51] while LRRC8C mutations affect cell volume regulation, contributing to various disorders [52]. VRAC permeability varies with subunit composition, influencing metabolite transport [53].

Thus, evaluating the permeability of other large-pore channels is essential for understanding their physiological and pathological roles.

## Conclusion

Our methodology offers a technically accessible approach for comparing the permeability of different Cx hemichannels by integrating electrophysiological and fluorescence imaging data. While our approach requires larger datasets than simultaneous measurement techniques, its experimental flexibility and scalability make it valuable tool for studying permeability of Cx hemichannels, and potentially, other types of large-pore channels. Future refinements, including improved diffusion model and machine learning applications, may further enhance utility of this methodology.

## Author Contribution

T.K. and M.S. conceived the study. T.K., L.Kr., V.K.V., and M.S. designed the research. T.K. and L.Kr. performed electrophysiological and fluorescent imaging experiments. L.Ke., A.K., and D.C. developed the AI-based cell segmentation system and performed its testing. M.S. developed the mathematical and statistical models. A.C. and M.S. conducted data analysis and simulations. T.K., V.K.V., and M.S. prepared the manuscript.

